# Increased resolution of African Swine Fever Virus genome patterns based on profile HMM protein domains

**DOI:** 10.1101/2020.01.12.903104

**Authors:** Charles Masembe, My V.T. Phan, David L. Robertson, Matthew Cotten

## Abstract

African Swine Fever Virus (ASFV) was originally described in Africa almost 100 years ago and is now spreading uncontrolled across Europe and Asia and threatening to destroy the domestic pork industry. Neither effective antiviral drugs nor a protective vaccine are currently available. Efforts to understand the basis for viral pathogenicity and the development of attenuated potential vaccine strains are complicated by the large and complex ASFV genome. We report here a novel method of documenting viral diversity based on profile Hidden Markov Model domains on a genome scale. The method can be used to infer genomic relationships independent of genome alignments and also reveal ASFV genome sequence differences that alter the presence of functional protein domains in the virus. We show that the method can quickly identify differences and shared patterns between virulent and attenuated ASFV strains and will be a useful tool for developing much-needed vaccines and antiviral agents to help control this virus. The tool is rapid to run and easy to implement, readily available as a simple Docker image.

## INTRODUCTION

African Swine Fever Virus (ASFV), belonged to the *Asfarviridae* family, was first described in Kenya nearly 100 years ago (1). The virus is endemic in most sub-Saharan African countries where it naturally infects warthogs and bush pigs and is frequently transmitted via soft ticks. In sub-Saharan Africa, infections of warthogs and bush pigs have a typically mild disease outcome. In domestic swine or wild boars, ASFV infections can result in a more serious disease with much greater mortality between 90 – 100%. Of great concern for animal welfare and the food industry, ASFV infections are responsible for increasing swine mortality in several parts of the world (2). Outside of Africa, the virus has been previously reported in Portugal, and in Haiti in sporadic outbreaks, probably as an import from West Africa (3)(4). Since the virus’s first European appearance in Georgia in 2007, the virus has spread in wild boar populations in Europe (reviewed in (5)), with currently 3,608 cases reported in wild boar and 1,413 cases in swine as of 1 June, 2019. Disturbingly high prevalence of ASFV has been found in Chinese dried pig blood used as porcine feed additives with all 21 tested samples testing positive by PCR as well as the generation of a full ASFV genome sequence (6). Furthermore, ASFV sequences have been identified in Chinese pork imported into Korea (7). These recent European and Asian incursions and outbreaks involve p72-Genotype II ASFV and appear not to involve the soft tick stage as originally observed in some parts in Africa. At the time of writing, neither antiviral drugs/agents nor an effective vaccine are available to stop the epidemic.

The ASFV virion is enveloped, spherical or pleomorphic in shape with a diameter of 175-215 nm. The virus has a linear, dsDNA genome of 170-195 kb with complementary terminal sequences. The ASFV genome encodes >150 open reading frames (ORFs) (8). In addition to known viral structural and replication proteins, there are a large number of ORFS with undefined functions. These include the multi-gene families (MGFs) that show frequent duplication, deletion or inversion across the virus family (8). Multiple examples of attenuated ASFV strains encoding changes in MGF content, indicate that MGFs have a role in ASFV virulence (9) (10) (11) (12) (13) (14) (15) (16) (17). However, the complexity of the MGF families and the nature of their sequence changes in ASFV evolution make it difficult to accurately ascribe specific changes in the ASFV genome to changes in phenotype. A simplified tool for monitoring these potentially functional changes would benefit the field and may aid in making a safe attenuated vaccine strain as well as to guide efforts to develop antiviral therapies.

The p72 gene (1,942 bp) is frequently used for PCR diagnosis of ASFV (18). Additional genes used for the diagnosis include the central variable region (CVR) of pB602L gene and p54 protein (encoded by E183L gene, an antigenic structural protein involved in the viral entry). Currently, there are 24 ASFV genotypes described based on p72 sequences (19), with the two most recent genotypes found in Ethiopia (20) and Mozambique (21). There have been efforts to classify ASFV strains, including using 3 ORFs (22) (23) (24) (25), the p72 gene (26), and the pB602L gene (27). In general, these methods have been limited to small portions of the ASFV genome (i.e. < 1% of the genome size), which are not likely to capture the full evolutionary history of the virus. Important drivers for this activity are efforts to understand the pathology of the virus infection, the components of a protective immune response and most important for vaccine development, the generation of attenuated but still immunogenic virus strains that may be used for vaccine applications. Altogether, this would help prevent and control the transmission of this virus across continents.

We have been developing the use of encoded protein domains as a classification tool for viral genomic sequence data, for example, applied to *Coronaviridae* genome sequences (28). A domain is a functional unit of a protein; different combinations of domains will give rise to different functional proteins. Instead of using differences in nucleotide or protein sequences to identify possible changes across sets of evolutionary related viral genomes, employing the domain classification would inform not only the genome changes but also potentially functional alterations of the virus genomes. All protein domains are well described in the Pfam collection, available at http://pfam.xfam.org. Novel instances of a domain and its relative distance to a reference domain can be rapidly identified in query sequences using the software HMMER-3 (29). HMMER (available at https://hmmer.org/) was developed by Eddy *et al* (29) to rapidly search a profile database for sequence homologs employing profile hidden Markov models (profile HMMs) probabilistic models. This strategy can be used to describe all domains encoded by a viral genome. A matrix of these domain scores can then be used to compare and cluster sets of ASFV genomes similar to a sequence-based phylogenetic analysis. We have developed these ideas further in this work to explore ASFV genome diversity and evolutionary relationships, to provide some functional clues for differences in viral genomes and to help identify viral elements associated with attenuation, virulence or transmissibility.

## MATERIALS AND METHODS

### ASFV Genome collection

All ASFV full genomes were retrieved from GenBank (5 April 2019) using the query: txid137992[Organism] AND 170000[SLEN]:200000[SLEN] yielding 48 complete genomes. Two genomes were identical MK333180 and a genome derived from dried blood products MK333181, only MK333180 was retained for a final set of 47 genomes. The GenBank entries and original literature were searched for country, date and original host (tick, warthog, wildboar or domestic pig) as well as any indication of virulence derived from the original literature. A summary of the 47 genomes used for the analysis is in Supplementary Table 1.

### Pfam-A domain content

The Pfam domains encoded by ASFV genomes were identified using hmmsearch function of HMMER-3.2.1 (29), searching against the most recent Pfam database (Pfam 32.0, September 2018, 17929 entries) (30) (31). For each genome in the collection, all ORFs ≥ 75 amino acids (aa) were collected from both reading strands and then examined for the presence of Pfam content. A domain hit was retained if the domain_i-Evalue was ≤ 0.0001. Details of each domain instance were gathered including the position in the query genome, the length, the domain_i-Evalue, and the bit-score.

### Custom profile HMMs for the MGFs

All ASFV encoded MGF protein coding sequences were retrieved from GenBank as follows. An initial query to the NCBI Nucleotide database was made to retrieve complete or nearly complete ASFV genomes (txid137992[Organism] AND 170000[SLEN]:200000[SLEN] NOT patent). From the “Send to” menu, the option “all coding sequences” was selected and these entries were retrieved to a fasta file. MGF entries were selected from the complete ASFV coding sequence file by sorting for the presence of the term “MGF” in the coding sequence ID with a simple python script. This yielded a set of 660 MGF entries.

When screened for Pfam content, 127 of the 660 protein coding sequences failed to return a domain hit (at a lenient domain_i-Evalue cutoff of 0.01). These were classified in GenBank as MGF_100 (38 entries); MGF_110 (9 entries); MGF_300 (39 entries); and MGF_360 (41 entries). To increase resolution for ASFV genome comparisons, profile HMMs were prepared for these proteins as follows. The 660 MGF ORFs were clustered using Usearch (32) at an aa fraction sequence identity of 0.75. Initially clustering pilots were performed at identities of 0.95, 0.90, 0.85, 0.80. 0.75, 0.70 and 0.65 (the lowest ID cut-off recommended for Usearch clustering). The 0.75 clustering gave the best separation of the coding regions into groups that corresponded to the GenBank annotation. In general, clustering followed the annotation, however several MGFs were further divided into subfamilies at this identity cut-off resulting in a set of 45 MGF subfamilies. Each MGF subfamily was aligned with Mafft (33), and a profile HMM was built using hmmbuild (29).These custom profile HMMs were used in combination with the identified Pfam profile HMMs (see Results).

The computational tools for performing this analysis are openly available as a platform independent Docker image of the tool and instructions for installing and using the tool have been made available (see Availability section and Readme document in the Supplementary Data). The Docker image contains the Unix, python, biopython SciKit and HMMER-3 modules need to run the classification, and the set of 511 HMMs (469 from Pfam plus 45 custom profileHMMs from MGF families) that were used to classify ASFV genomes. Outputs from the classification tool are a clustermap showing the relationship between the genomes, and a CSV table listing all domains identified in each genome, their position, length and coding strand in the genome and a flag indication high or low variance. This CSV table is useful for investigators wishing to explore the identified domains further or to investigate differences between genomes.

### UK domain analysis

The UK protein coding sequence was retrieved from the GenBank entry NC_001659 for the BA71V strain and used in an online blast search (megablast default settings) to identify closely related sequences. Using the download menu, all hits (39 entries, 1 October 2019) were retrieved to a fasta file, the UK domain coding sequence from the NC_001659 genome was added, and the set was translated into protein sequences using Geneious, aligned in Mafft (33) (mafft --auto --preservecase ASFV_UKorf_set_aa.fas > ASFV_UKorf_set_aa_aln.fas) and Geneious was used to calculate pairwise aa differences and to visualize protein changes across the alignment. The Pfam domain content of the UK protein coding sequence set was determined as described above, identifying only the UK domain at a domain_i-Evalue cutoff of < 0.0001. The domain bit-scores were collected for the set and compared to the pairwise aa differences (see Supplementary Figure 1).

The 47 ASFV full genome sequences available in GenBank were aligned using Mafft (33) and manually checked in AliView (34). Maximum-likelihood (ML) phylogenetic tree of the p72 gene was constructed in RAxML (35) under the GTRGAMMA model of substitutions and bootstrapped for 100 pseudo-replicates. The tree was mid-point rooted for clarity and branches were drawn to the scale of nucleotide substitutions per site, and bootstrap values > 75% are indicated.

## RESULTS

### Documenting Pfam content of ASFV

Initially, we identified all profile HMM domains from the Pfam collection that were encoded in a set of 47 ASFV genomes. Using a domain_i-Evalue cutoff of 0.0001 (a measure of the probability of finding the domain by chance), 82 domains were identified at least once in the set of 47 genomes, and 17 domains were found twice or more in the set indicating repeat occurrences in some genomes (see Supplementary Table 2). The domain content and their scores (from Pfam plus custom MGF domains) were then used to examine patterns of the 47 ASFV genomes in GenBank in the following manner. Briefly, for each genome a total score for each domain was generated by summing the individual domain scores (taking into account multiple instances of the same domain). For each domain column in the matrix, the scores were normalized by dividing each value by the maximum value; domains that showed > 0.03 variance in their score across the set of 47 genomes were retained and used for hierarchical clustering. A schematic presentation of the process is shown in Figure 1.

**Figure 1.**
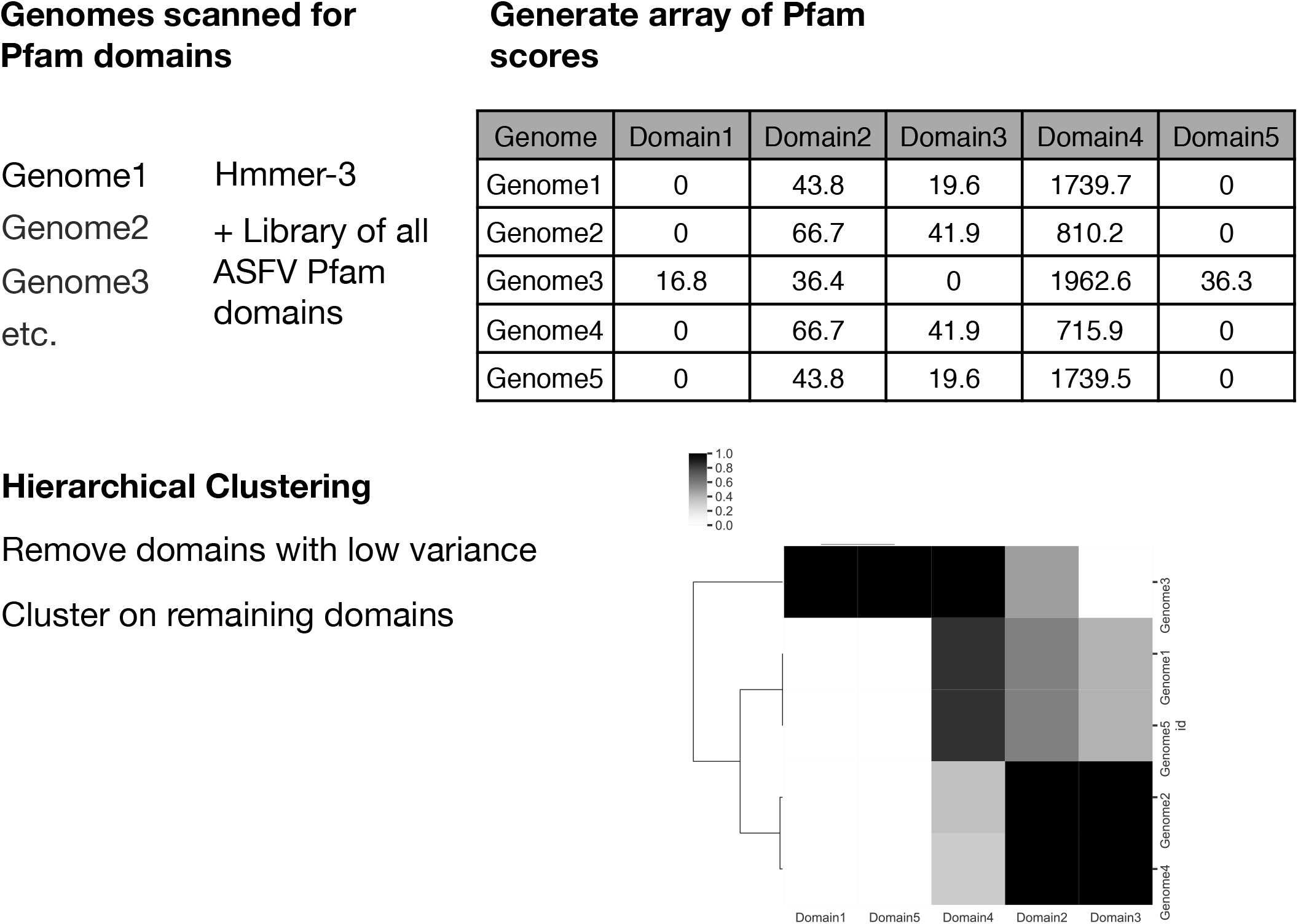
The process of genome clustering with HMMs. Each full ASFV genome was scanned for Pfam and MGF domain content (step 1), the domain scores were collected, built into a matrix and normalized to fraction of highest score in the set (step 2). Domains with low variance across the entire set were removed and hierarchical clustering of the genomes was performed using the high variance domains (step 3).

### Domain variability measured by this method

As an illustration of the domain-classification approach, we examined the UK gene’s ORF encoding a 96 aa protein expressed early in ASFV infection (36). Although the protein is nonessential for growth in porcine macrophage cell cultures, deletion of the UK coding region reduces the virulence of ASFV in domestic pigs (36). A set of ASFV “UK” coding regions was retrieved from GenBank, an alignment of the proteins set is shown in Supplementary Figure 1A, revealing 22 aa differences between the most divergent forms of the protein. Following the HMMER-2 search of the UK ORFs, the Pfam domain score (bit-score) for the UK domain varies across the set with a bit-score value of 227.7 for perfect match. In support of the use of this metric, there is a highly significant negative correlation between Pfam domain score with the pairwise aa distance (Supplementary Figure 1B). Of note, the Pfam UK domain entry was constructed using the ASFV reference strain NC_001659 UK protein as a model and the HMMER-3 score is correlated with the differences of query domains from this early ASFV sequence. Thus, a HMMER-3 search can be used both to find members of a domain family in a query genome as well as to provide a quantitative score (bit-score) of the distance of the query domain from the model domain.

### Documenting Pfam content of ASFV

We identified all profile HMM domains from the Pfam collection that were encoded in a set of 47 ASFV genomes. Using a domain_i-Evalue cutoff of 0.0001 (a measure of the probability of finding the domain by chance), 82 domains were identified at least once in the set of 47 genomes and 17 domains were found twice or more in the set indicating repeat occurrences in some genomes (see Supplementary Table 2). As described above, the domain content and their scores (from Pfam plus custom MGF domains) were then used to examine patterns of the 47 ASFV genomes in GenBank.

The 47 full ASFV genomes were ordered by hierarchical clustering based on the Pfam + MGF domain scores and compared to a p72 ML tree with the major genotypes in each analysis indicated by colored boxes (Figure 2). In validation of our approach, the domain-clustering (Figure 2, panel B) groups genomes in nearly the same pattern as p72 ML tree topology (Figure 2, panel A), which is a current standard practice to genotype ASFV strains. Differences include the phylogenetic position of older genomes and those genomes obtained from tick samples. Of note, the genotype II (GII) viruses, that are spreading globally, clustered into a monophyletic group on the p72 ML tree (green shaded, Figure 2A). Interestingly, the domain clustering showed that the Estonian genome (GenBank LS478113, identified from a wildboar in 2014 (37)) possesses a large 14kb deletion, lacking functional domains MGF_110 1L-12L compared to other genotype II ASFV viruses (Figure 2B). Additionally, within the GII ASFV viruses, strains FR682468 and MH766894 show changes in the DUF4509 domain (associated with MGF_360 genes). In addition to diversity in the MGF domains, there is diversity (with variance ≥ 0.03) in the 11 domains (AAA_22, Ank_2, Ank_5, ATPase_2, mRNA_cap_enzyme, Nodulin_late, P12,RIO1, SHS2_Rpb7-N, TFIIS_M, UK) observed across different genotypes. None of these domain absence/presence are revealed from a p72 ML tree (Figure 2A) that is typically used to genotype these viruses.

**Figure 2.**
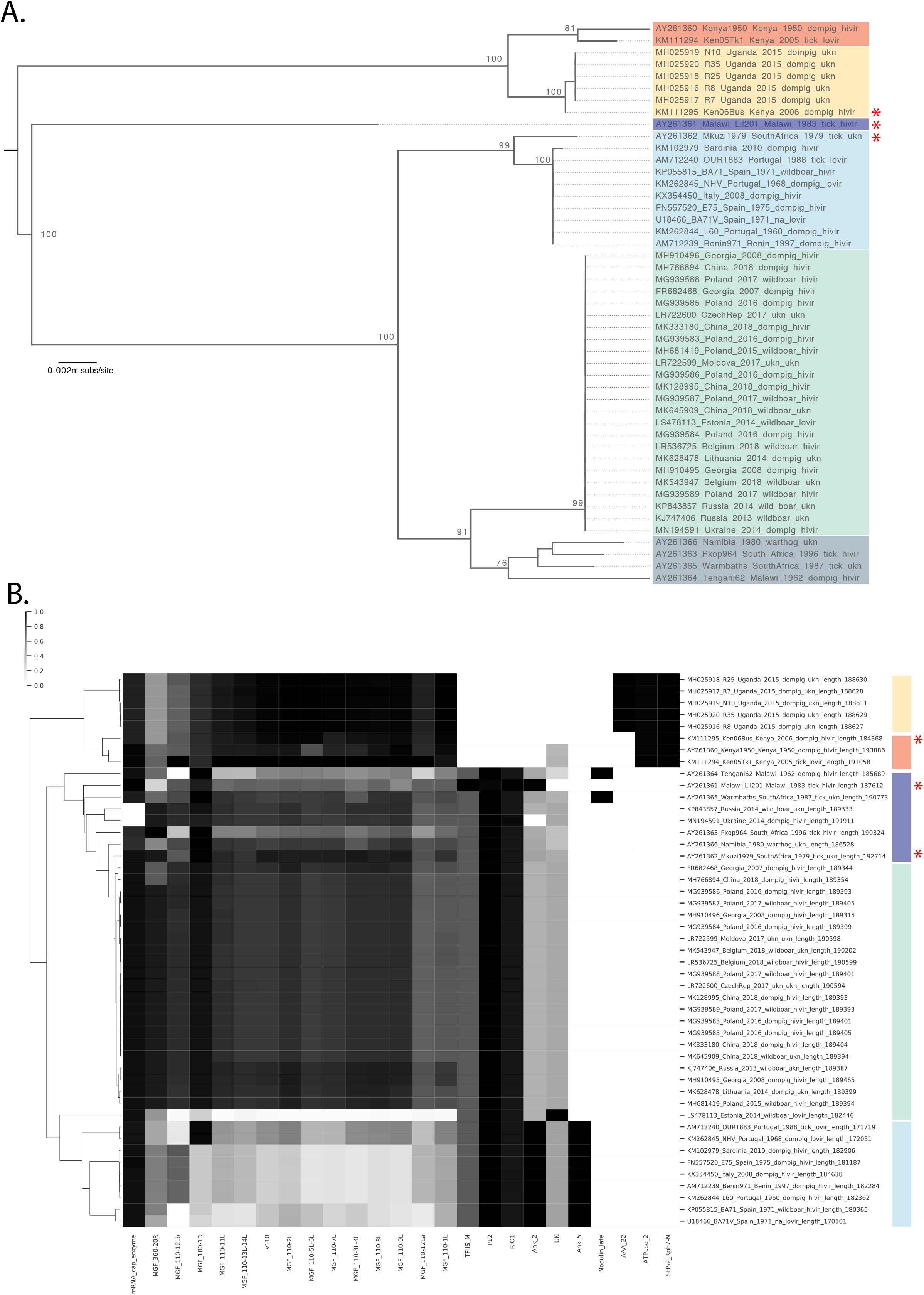
Panel A. The p72 maximum-likelihood phylogenetic tree. The coding sequences of p72 gene from the 47 ASFV genomes available in GenBank were aligned in Aliview. ML tree was inferred using RAxML under GTRGAMMA model of substitutions with 100 bootstraps (see Methods for further details). The tree was mid-point rooted for clarity and branches were drawn to the scale of nucleotide substitutions per site (indicated in nucleotide substitutions/site), and bootstrap values ≥75% are indicated. Genotypes are indicated by colored boxes, with the Genotype II in green. **Panel B. The domain cluster-map classification of 47 ASFV genomes.** The 47 ASFV genomes were examined by their Pfam content (see Methods). The bit-scores for all domains identified with domain_i-Evalue ≤ 0.0001 were collected for each domain, a matrix was prepared and subjected to hierarchical clustering (see Materials and Methods) based on domain whose normalized values showed ≥ 0.03 variance. In both panels, the genotypes are indicated with colored boxes. Genome IDs shown on node labels (panel A) and Y axis (panel B) include GenBank accession number, strain name, country, date, host, virulence and length in nucleotides. For both panels, genomes with incongruent placement between the two methods are highlighted with a red asterisk.

### Domains associated with Multigene Families (MGFs)

Five MGFs have been defined (MGF 100, 110, 300, 360 and 505/530) with the naming based on the mean number of amino acids in the gene product. All annotated ORFs from 47 complete genome entries in GenBank were collected (660 total entries, MGF_100: 38; MGF_110: 148; MGF_300: 46; MGF_360: 267; MGF_505: 160 entries) and examined for Pfam domains. Three MGFs consistently encoded at least one domain (i.e. all members of that MGF family were found to encode a particular domain). These were MGF_110: domain v110, MGF_360: domain ASFV_360, MGF_505: domain DUF249. To capture the diversity in these MGFs, we prepared individual profile HMMs from a comprehensive set of MGF ORFs. Briefly, we grouped each MGF protein by aa sequence identity and identified 45 MGF subfamilies and then constructed custom profile HMMs for each of these (see Methods). We then analyzed the clustering pattern of all MGF ORFs based on their custom profile HMMs (Figure 3). Most MGFs clustered within their annotated family, evidenced by the rectangle of shared score similarities surrounding the large clusters of MGF_100 and MGF-110, MGF_360, MGF_505 (Figure 3). However, a subset of 10 MGFs appeared different from the main MGF group bearing their name (Figure 3, red boxes, IDs with asterisks). For example, several ORFs annotated as MGF_505-11L have less than 0.85 aa sequence identity (fractional identity (32)) with other MGF_505 family member and their domain scores cluster them to a unique sector of the graph (Figure 3 red box). There is a similar pattern for MGF_360-15R, MGF_300-1L and 2R, MGF_360-18R, MGF_300-4L and MGF110-12L revealing greater domain/functional variety in these genes than previously appreciated.

**Figure 3.**
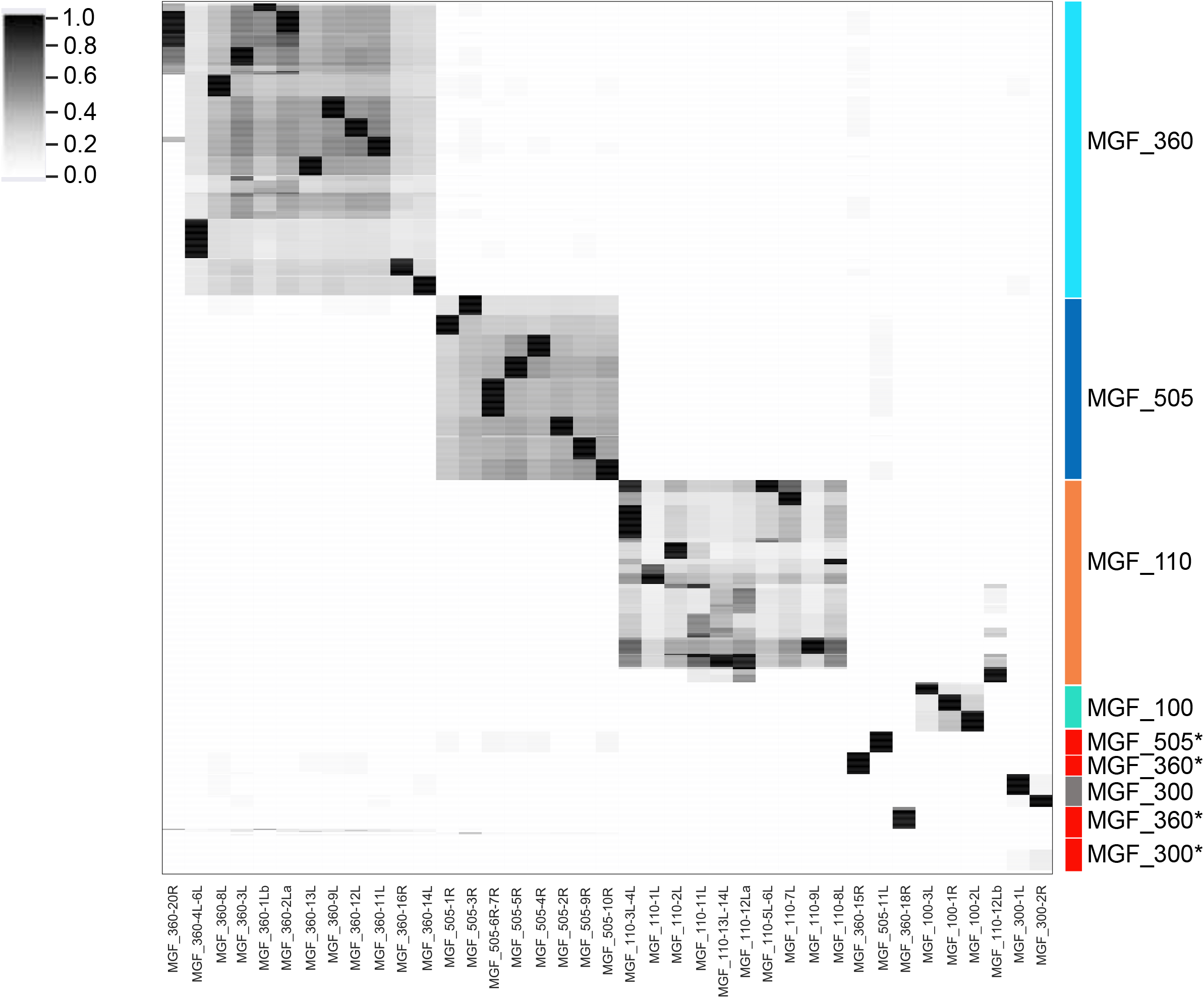
Hierarchical clustering of all available ASFV MGF protein sequences. All available ASFV MGF proteins (N=660) were retrieved from GenBank, clustered at an amino acid fractional identity 0.85 and a profile HMM was prepared from each of the 45 alignments (ASFV_HMM45) using HMMER3 (29). The same set of 659 proteins were then examined for ASFV_HMM45 content at an domain_i-Evalue threshold of 0.0001, bit-scores were collected and used to prepare a matrix describing the set of proteins. The matrix was then subjected to hierarchical clustering and a clustermap prepared. Each column represents one of the 45 profile HMMs, each row represents an MGF protein. Major clusters are indicated to the right, unconventional domains that do not cluster with other members bearing the same GenBank MGF family annotation are marked in the red box.

### Changes in domain copy number

It has been previously noted that MGF counts vary with ASFV genotype and also between attenuated and virulent strains. This is illustrated in Figure 4 where we have plotted specific domain counts by sample date and virus genotype. As clearly shown in Figure 4, viruses of genotypes GII and GIX possess higher levels of MGF_110 and MGF_360 specific domains. A few domains were observed to be absent from GII and GIX genomes, for example an Ankyrin 4 domain found in some genotypes is not present in GII or GIX (Figure 4).

**Figure 4.**
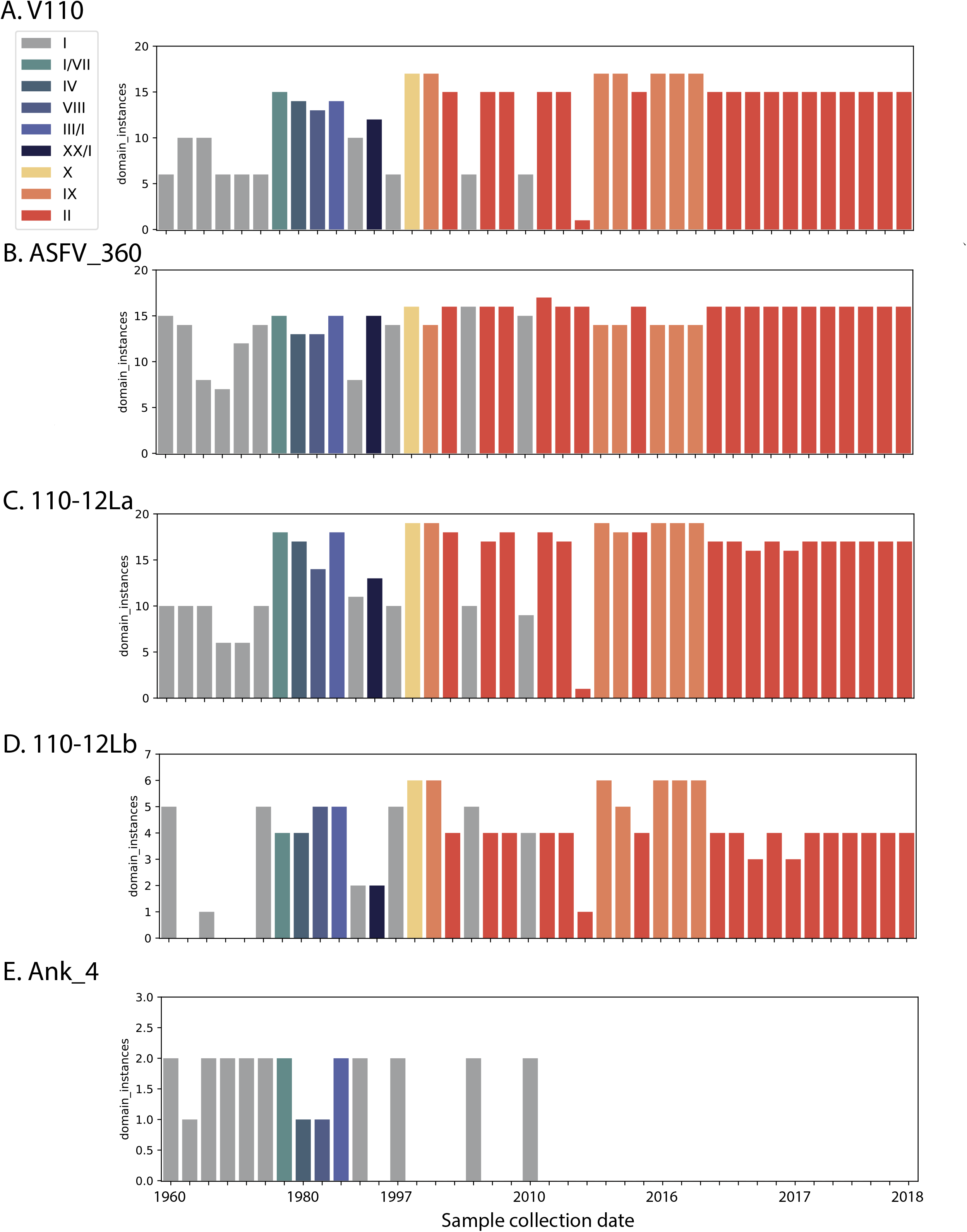
Changes in domain copy numbers. The total number of domains detected per genome was plotted per genome, organized by sample date and coloured by ASFV genotype (see legend insert for color code). Domains examined are panel A: Pfam v110 domain (found on MGS_110 family members), panel B: Pfam ASFV_360 domain (found on MGS_360 family members), panel C: the custom domain MGF_110-12La, panel D: The custom domain MGF_110-12Lb and panel E: the Pfam doman Ank_4. Genome ids (X axis) include Genbank accession number, strain_name, country, date, host, virulence and length in nucleotides.

Of potential importance to disease status, it has been observed in several analyses that changes in MGF numbers might result in altered viral properties. A deletion of a large 5’ region including multiple MGF_110 elements was associated with attenuation of an Estonian ASFV strain (37). Two GI viruses Lisboa60 (L60, KM262844, a virulent strain) and NH/P68 (NHV, KM262845, a non-virulent strain) studied for their differences in virulence revealed differences in 4 MGF families (MGF_100, MGF_110, MGF_360, MGF_505 (38). The attenuated strain NHV showed increases in MGF_100 and MGF-110 scores and decreases in MGF_360, MGF_505 scores. MGF_110-12La, an unconventional MGF_110 family member, has higher domain counts in GII strains (Figure 4, Panel C), while MGF_110-12Lb, an unconventional MGF_110 family member, has the highest domain counts in GIX Uganda viruses (Figure 4, Panel D). The Ank-4 domain is not detected in GII, GIX viruses. Ankyrin motifs are typically found in scaffolding and signaling molecules.

### Analyses of paired viruses

Finally, we applied the genome-scale domain comparison method to examine pairs of ASFV strains with reported differences in virulence. Such analyses are crucial in efforts to understand the molecular basis for attenuation or virulence and to guide efforts for vaccine design.

For example, a naturally-occurring ASFV variant was recently described from Estonia that displayed attenuation in animal tests (37). The original report noted that the Estonian variant was missing 26 genes including 13 members of the MGF_110 family, 3 members of the MGF_360 family, deletions of MGF_100_1R, L83L, L60L and KP177R as well as a duplication and rearrangements (37). We applied the domain classification tool to compare the variant Estonian strain to contemporary viruses from Georgia, changes in protein domains are shown in Figure 5A with domains showing variation across the set of four related genomes indicated by changes in the cluster map. The MGF_110 and MGF_360 changes previously noted are clearly visible with reduced signals for these two families of genes (Figure 5A). Additional domain changes were observed including variations in the DUF4509, UK, PP1c_bdg and ASFV_L11L domains. The DUF4509 domain is found on a subset of MGF_360 domains and is consistent with the reported MGF_360 changes. The PP1c_bdg domain is found on a Phosphatase-1 catalytic subunit binding region that may influence apoptosis (39) and may be relevant for ASFV virulence. The ASFV_L11L domain also shows changes, this domain is found on the L11L gene which although reported to be non-essential for virus growth (40) was previously noted to be missing from attenuated viruses (37).

**Figure 5.**
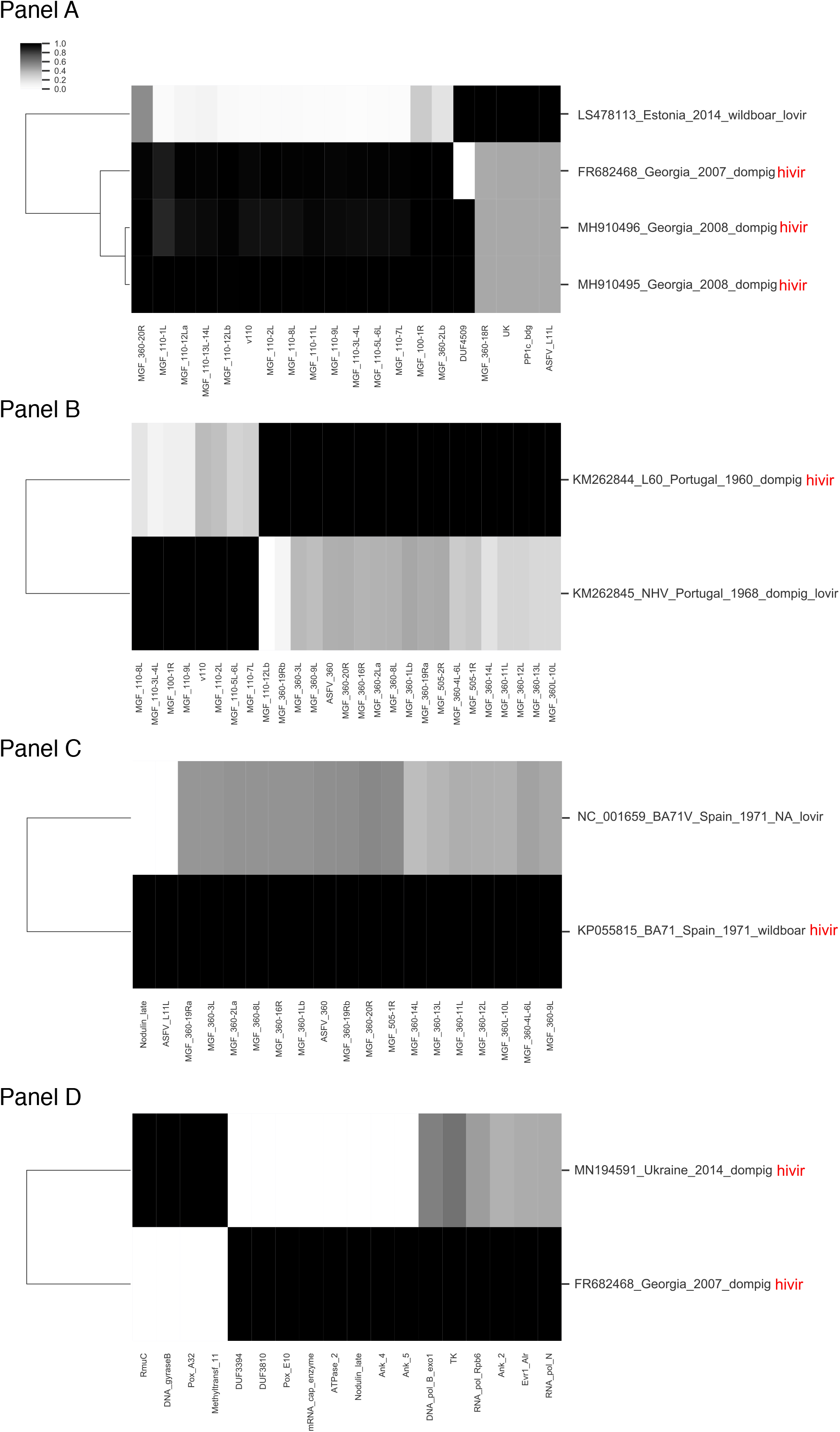
Differences in domains between paired ASFV strains. For each panel, the indicated genomes were examined for Pfam and MGF domain content, the bit-scores for all domains identified with domain_i-Evalue ≤ 0.0001 were collected for each domain, a matrix was prepared and subjected to hierarchical clustering (see Materials and Methods) based on domain whose normalized values showed ≥ 0.03 variance. Genome ids (Y axis) include GenBank accession number, strain_name, country, date, host and virulence (lovir = low reported virulence, hivir = high reported virulence).

Other examples include the Lisboa60 (L60) virulent strain and the NH/P68 (NHV) non-virulent strain, which have been described and compared for virulence differences (38). Domain differences between the two strains confirms the previously reported changes in MGFs (100, 110, 360 and 505, Figure 5B). Also, BA71 and BA71V are a pair of virulent/attenuated ASFV strains. The BA71V strain was adapted to cell culture and showed attenuation accompanied by the loss of MGF_360 and 505 genes (41) (42). The domain differences between the two strains are consistent with previously reported differences in the MGF_360 and MGF_505 genes. In addition, the ASFV_L11L domain and a Nodulin_late domain show a change in signal in the attenuated strain (Figure 5C). The observed changes in ASFV_L11L in two quite different pairs of virulent/avirulent ASFV strains is notable and the role of the ASFV_L11L membrane protein should be re-examined in more detail.

## DISCUSSION

We have demonstrated the utility of a novel method of characterizing ASFV-encoded protein diversity on a genome-scale based on profile Hidden Markov Model descriptions of conserved protein domains. The method exploits the Pfam collection of profile HMMs (43) as well as the rapid and sensitive HMMER3 software (29). The standard methods of accurately comparing large virus genomes requires the careful preparation of a full-length genome alignment of the ~190 kb ASFV genome combined with a maximum likelihood phylogenetic tree inference coupled with bootstrapping to check cluster reliability. The combined phylogenetic analysis might take several days to complete and is further complicated by the large size and frequent gene deletions and duplications in the ASFV genome making an accurate and reproducible alignment quite difficult to generate. In comparison, the domain method described here requires no alignment and can be performed from an unaligned fasta file of the genome sequences through to hierarchical clustering in minutes. The clustermap analyses reported for 47 ASFV full genomes was performed in approximately 3 minutes run-time on a standard laptop (in this case a 2018 MacBook Pro with 2.7 GHz Intel Core i7, and 16 GB of memory). The method will be useful for quality control of newly assembled genomes and for exploring novel ASFV genomes as they are sequenced and annotated, as well as for comparing genomes with varied clinical, epidemiological and phenotypic outcomes. The combination of our approaches with the viral outcomes are important in efforts to develop an effective and safe ASFV vaccine.

We have identified greater diversity in the 5 MGF families than previously noted. We further reveal the presence of a set of unconventional MGFs (Figure 3) that appear distinct to ASFV. Their presence and evolution will need to be monitored in future studies. Indeed, the process of MGF evolution may be an important part of ASFV evolution and the current work provides novel tools for monitoring changes in these possibly high consequence genes. Grouping MGF genes in only 5 categories may result in a loss of information, obscuring important details necessary for understanding ASFV transmission, virulence and attenuation.

The domain method described here also allows a rapid assessment in both the qualitive features of encoded domains, and a reported a bit-score for each identified domain, which is a protein distance from the model domain. Furthermore, the method also reports copy number changes in domains. For example, examining changes in domain instances showed that the GII ASFV strains, responsible for large global outbreak of ASF, encoded a substantial increase in several MGF gene families (Figure 4). These changes may be an important part of the replication success of the virus and warrant further investigation.

The added benefit of domain-based classification is its alignment-free feature. The resolution of any phylogenetic constructions relies heavily on accurate alignment of homologous regions of sequences. In the case of ASFV, there are differences in MGFs across different ASFV strains, either duplications or deletions, which are very difficult and time-consuming to reliably align. Furthermore, if certain genes are missing from some of the genomes for some of the alignment, this region of the alignment may be masked in the entire alignment and will not contribute to the phylogenetic signal. However, such deletions, duplications or inversions of domains are captured by the domain scoring system used and may be an important component of the increased resolution of the domain method.

In conclusion, hierarchical clustering based on profile HMM domain scores has provided a rapid method of comparing similar genomes to identify differences in the encoded proteins. We applied the method to three sets of ASFV genomes from contemporary outbreaks with known phenotype differences in their ability to replicate in and kill pigs (Figure 5). The novel method identified previously noted differences (primarily in the encoded MGF genes) but revealed an additional set of changes that should be further explored as potential virulence factors. These functions may be important to remove or alter in efforts to generate attenuated yet immunogenic viruses.

Finally, we note that the computational tools for performing this analysis are openly available as a platform independent Docker image of the tool and instructions for installing and using the tool have been made available. We hope that by providing these computational methods as easy to implement tools they may help contribute to efforts to control this virus.

## AVAILABILITY

The computational tools for performing this analysis can be downloaded as a platform independent Docker image using this command (docker pull matthewcotten/asfv_class_tool). Instructions for installing and using the tool are available in the Supplementary Data Readme file.

---

**Supplementary Figure 1.**
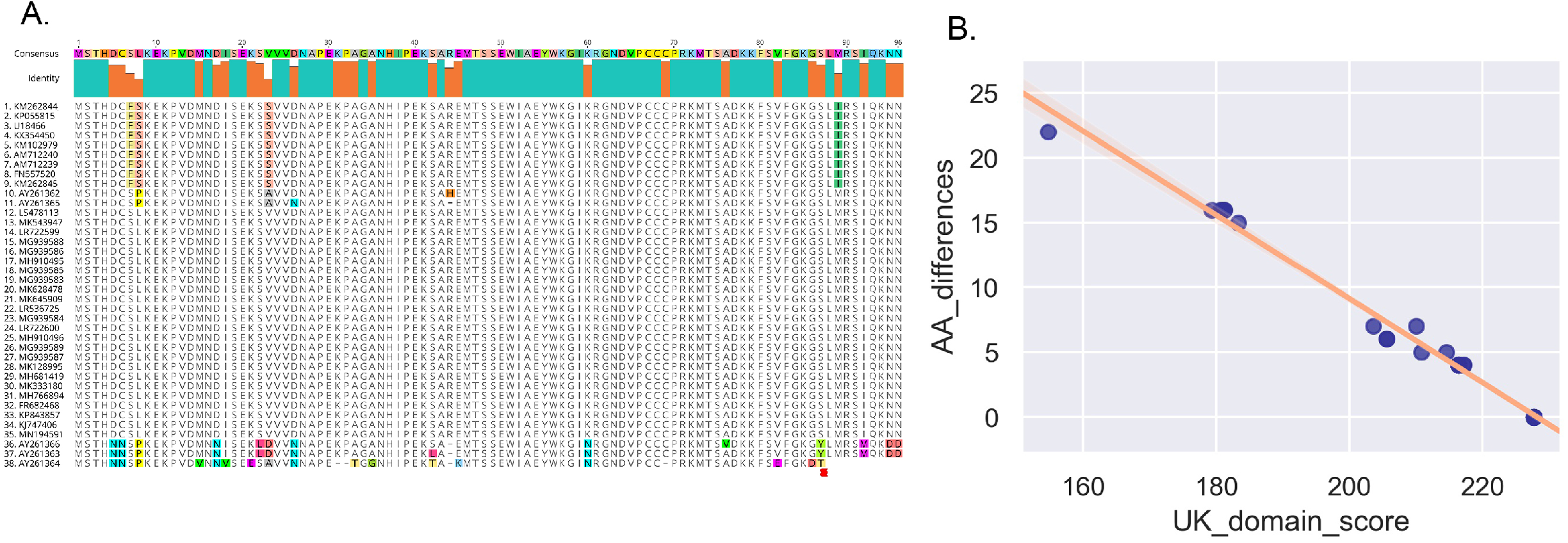
**Panel A**. All available ASFV “UK” ORF sequences from Genbank full genomes were translated into protein sequences, aligned in Mafft (33) and differences in the sequences relative to the consensus were visualized using Geneious (see Methods for details). **Panel B.** HMMR3 was used to screen the protein set for Pfam profile HMMs, the UK domain was detected and the bit-score for the domain from each sequence was plotted as a function of the pairwise protein sequence distance from the consensus. The Pearson correlation coefficient for the two sets of measurements was −0.995.

**Supplementary Table 1.**
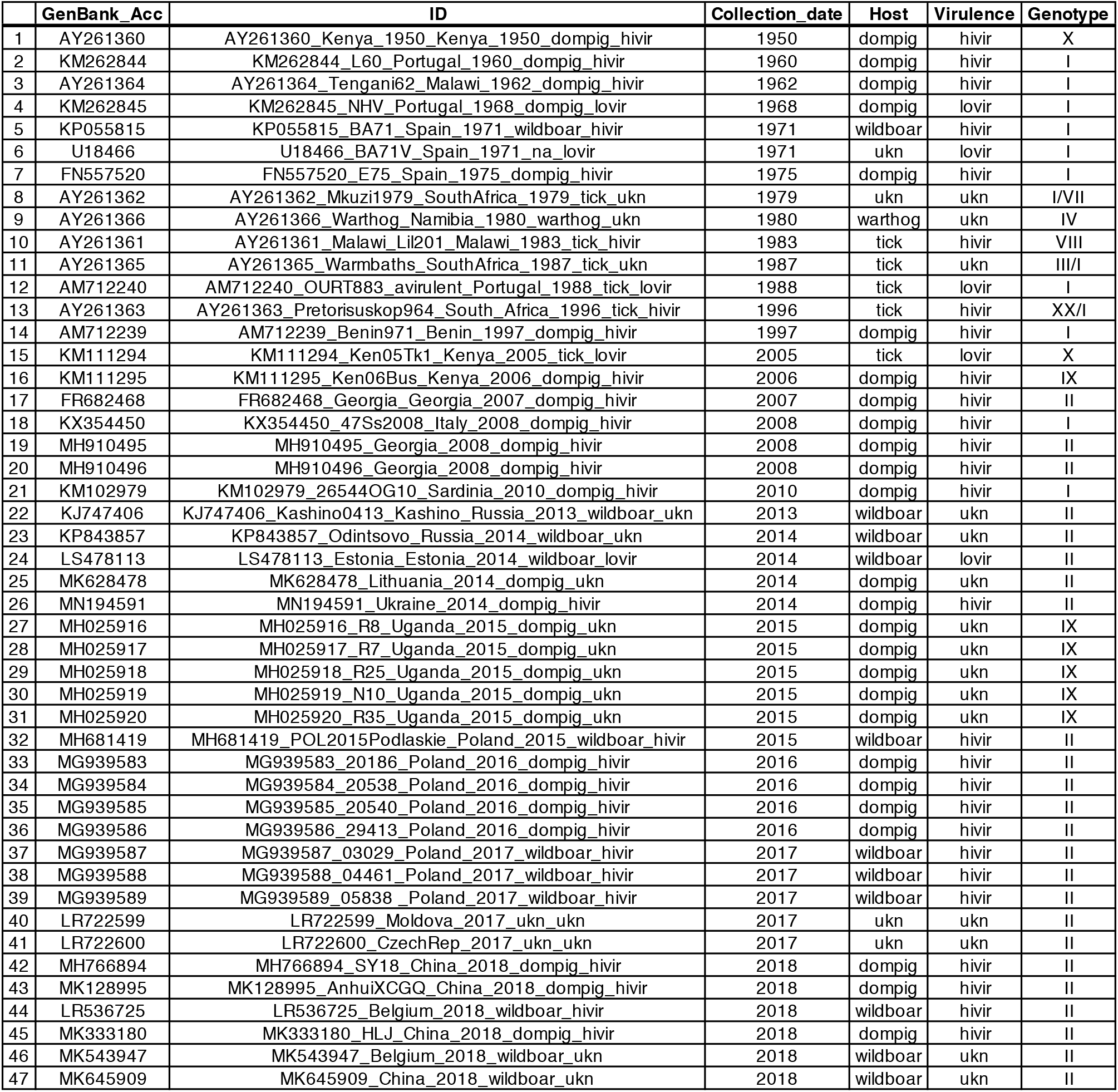

**Supplementary_Table_2.**
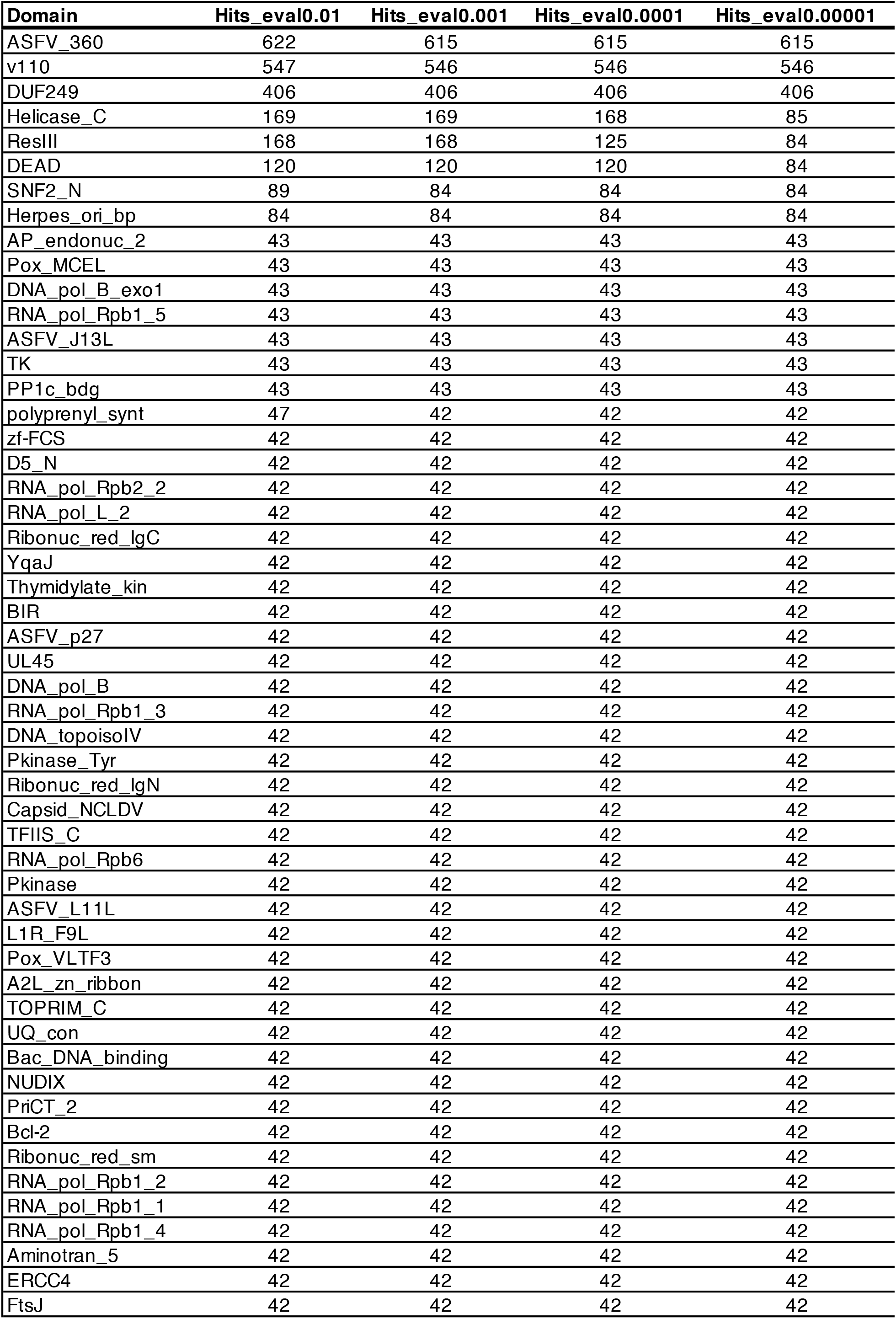
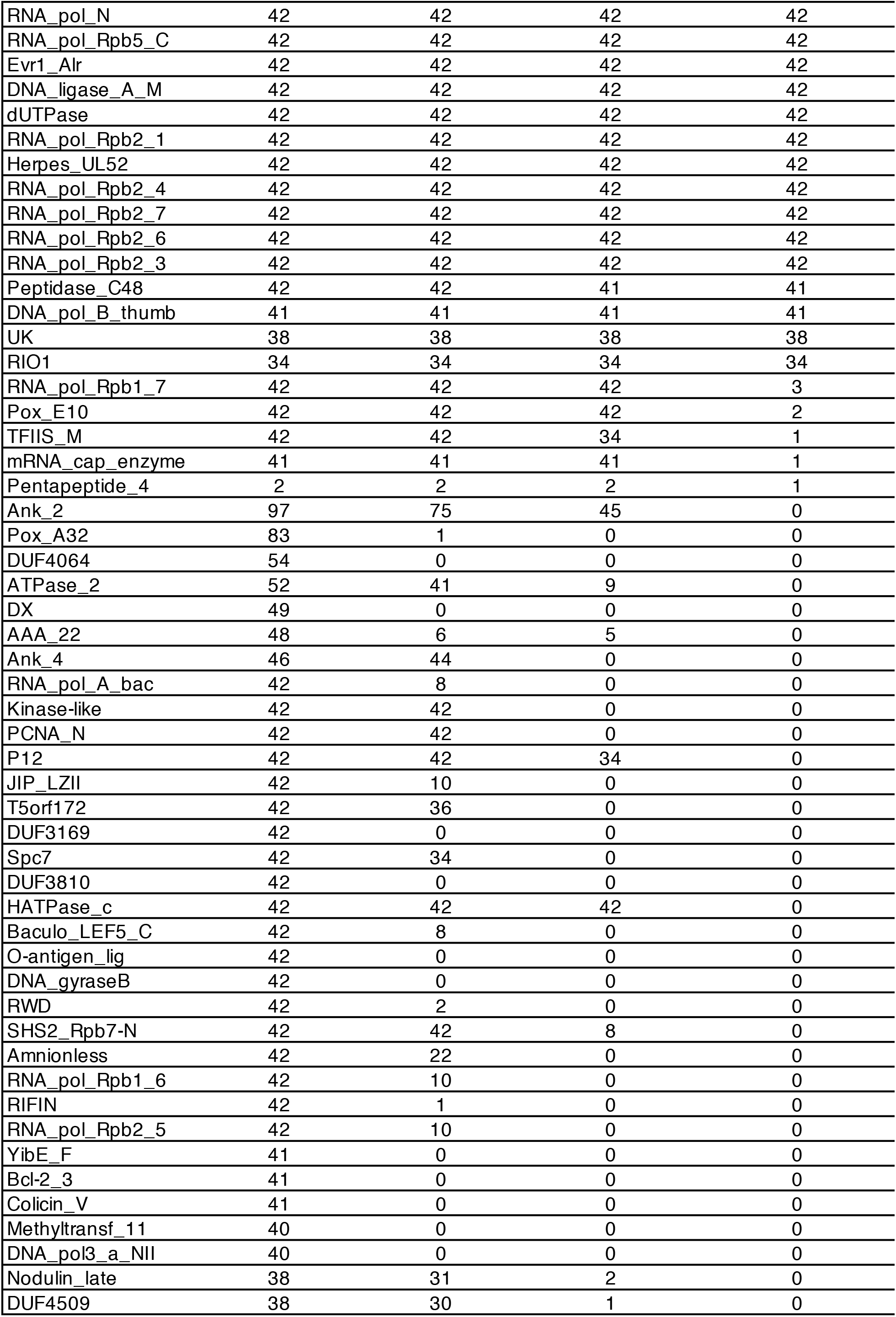
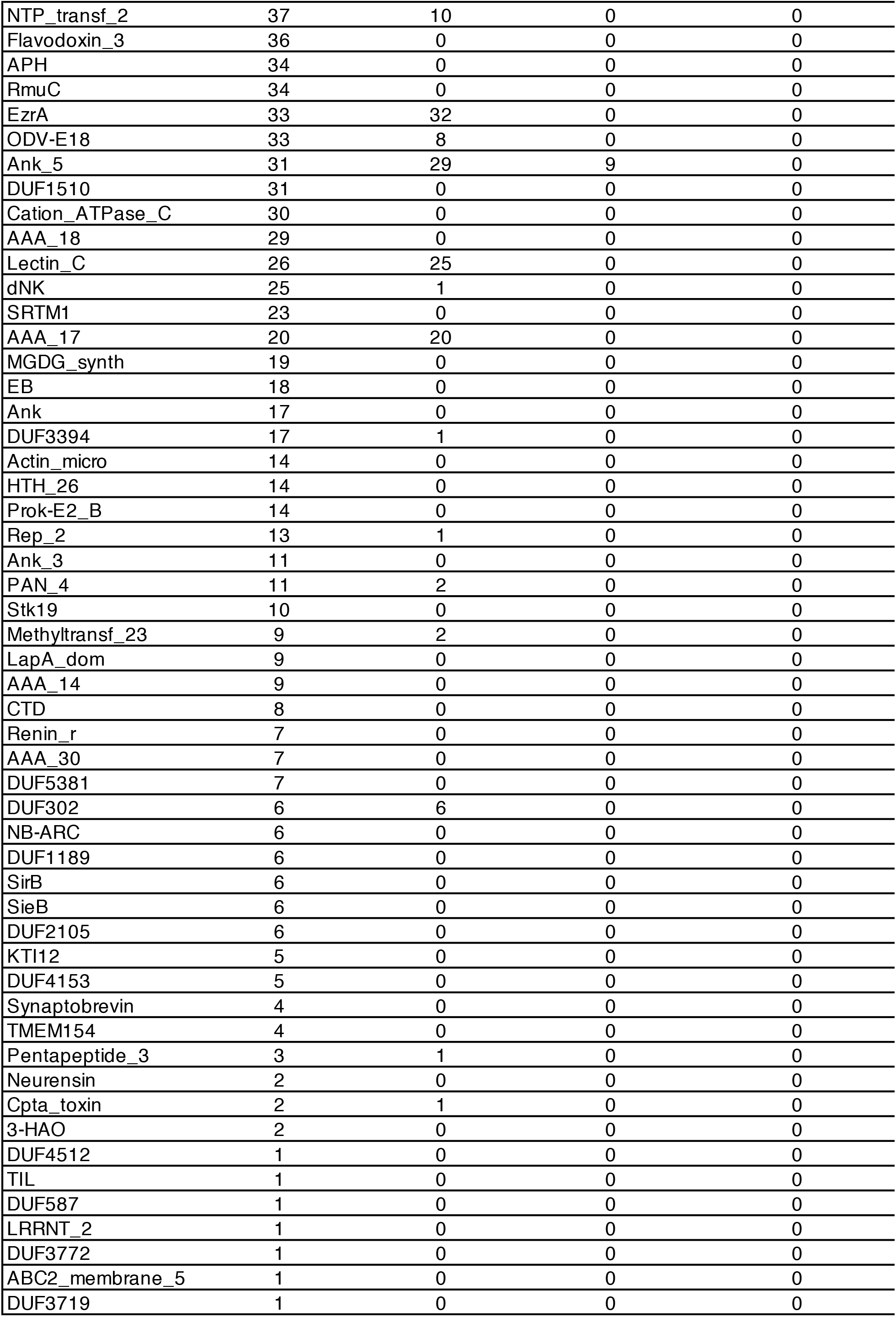
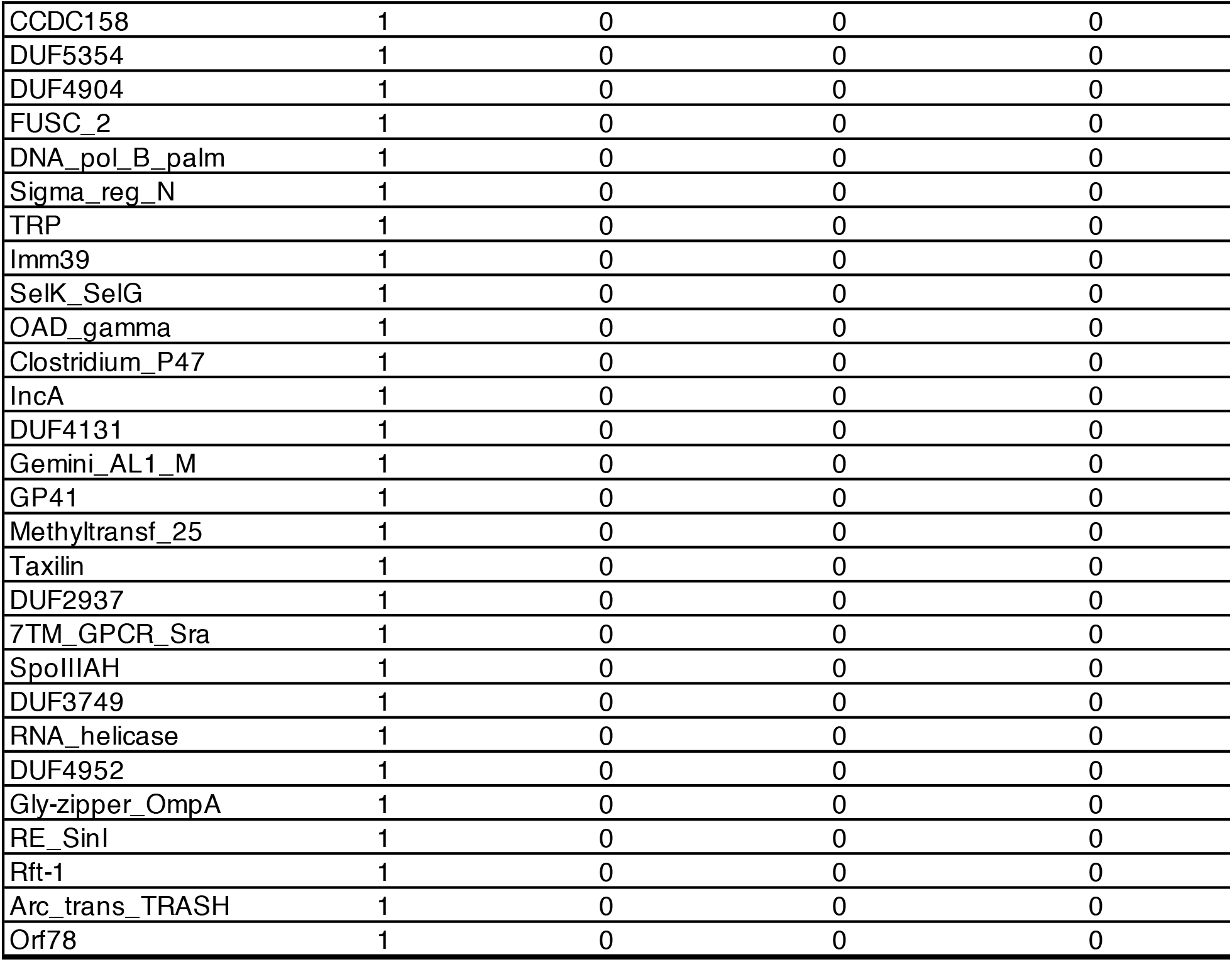

| Domain | Hits_eval0.01 | Hits_eval0.001 | Hits_eval0.0001 | Hits_eval0.00001 |
| --- | --- | --- | --- | --- |
| ASFV_360 | 622 | 615 | 615 | 615 |
| v110 | 547 | 546 | 546 | 546 |
| DUF249 | 406 | 406 | 406 | 406 |
| Helicase_C | 169 | 169 | 168 | 85 |
| ResIII | 168 | 168 | 125 | 84 |
| DEAD | 120 | 120 | 120 | 84 |
| SNF2_N | 89 | 84 | 84 | 84 |
| Herpes_ori_bp | 84 | 84 | 84 | 84 |
| AP_endonuc_2 | 43 | 43 | 43 | 43 |
| Pox_MCEL | 43 | 43 | 43 | 43 |
| DNA_pol_B_exo1 | 43 | 43 | 43 | 43 |
| RNA_pol_Rpb1_5 | 43 | 43 | 43 | 43 |
| ASFV_J13L | 43 | 43 | 43 | 43 |
| TK | 43 | 43 | 43 | 43 |
| PP1c_bdg | 43 | 43 | 43 | 43 |
| polyprenyl_synt | 47 | 42 | 42 | 42 |
| zf-FCS | 42 | 42 | 42 | 42 |
| D5_N | 42 | 42 | 42 | 42 |
| RNA_pol_Rpb2_2 | 42 | 42 | 42 | 42 |
| RNA_pol_L_2 | 42 | 42 | 42 | 42 |
| Ribonuc_red_lgC | 42 | 42 | 42 | 42 |
| YqaJ | 42 | 42 | 42 | 42 |
| Thymidylate_kin | 42 | 42 | 42 | 42 |
| BIR | 42 | 42 | 42 | 42 |
| ASFV_p27 | 42 | 42 | 42 | 42 |
| UL45 | 42 | 42 | 42 | 42 |
| DNA_pol_B | 42 | 42 | 42 | 42 |
| RNA_pol_Rpb1_3 | 42 | 42 | 42 | 42 |
| DNA_topoisolV | 42 | 42 | 42 | 42 |
| Pkinase_Tyr | 42 | 42 | 42 | 42 |
| Ribonuc_red_lgN | 42 | 42 | 42 | 42 |
| Capsid_NCLDV | 42 | 42 | 42 | 42 |
| TFIIS_C | 42 | 42 | 42 | 42 |
| RNA_pol_Rpb6 | 42 | 42 | 42 | 42 |
| Pkinase | 42 | 42 | 42 | 42 |
| ASFV_L11L | 42 | 42 | 42 | 42 |
| L1R_F9L | 42 | 42 | 42 | 42 |
| Pox_VLTF3 | 42 | 42 | 42 | 42 |
| A2L_zn_ribbon | 42 | 42 | 42 | 42 |
| TOPRIM_C | 42 | 42 | 42 | 42 |
| UQ_con | 42 | 42 | 42 | 42 |
| Bac_DNA_binding | 42 | 42 | 42 | 42 |
| NUDIX | 42 | 42 | 42 | 42 |
| PriCT_2 | 42 | 42 | 42 | 42 |
| Bcl-2 | 42 | 42 | 42 | 42 |
| Ribonuc_red_sm | 42 | 42 | 42 | 42 |
| RNA_pol_Rpb1_2 | 42 | 42 | 42 | 42 |
| RNA_pol_Rpb1_1 | 42 | 42 | 42 | 42 |
| RNA_pol_Rpb1_4 | 42 | 42 | 42 | 42 |
| Aminotran_5 | 42 | 42 | 42 | 42 |
| ERCC4 | 42 | 42 | 42 | 42 |
| FtsJ | 42 | 42 | 42 | 42 |
| Supplementary\_Table\_2\_ASFV\_PFAM\_domains, page 2 |  |  |  |  |
| RNA_pol_N | 42 | 42 | 42 | 42 |
| RNA_pol_Rpb5_C | 42 | 42 | 42 | 42 |
| Evr1_Alr | 42 | 42 | 42 | 42 |
| DNA_ligase_A_M | 42 | 42 | 42 | 42 |
| dUTPase | 42 | 42 | 42 | 42 |
| RNA_pol_Rpb2_1 | 42 | 42 | 42 | 42 |
| Herpes_UL52 | 42 | 42 | 42 | 42 |
| RNA_pol_Rpb2_4 | 42 | 42 | 42 | 42 |
| RNA_pol_Rpb2_7 | 42 | 42 | 42 | 42 |
| RNA_pol_Rpb2_6 | 42 | 42 | 42 | 42 |
| RNA_pol_Rpb2_3 | 42 | 42 | 42 | 42 |
| Peptidase_C48 | 42 | 42 | 41 | 41 |
| DNA_pol_B_thumb | 41 | 41 | 41 | 41 |
| UK | 38 | 38 | 38 | 38 |
| RIO1 | 34 | 34 | 34 | 34 |
| RNA_pol_Rpb1_7 | 42 | 42 | 42 | 3 |
| Pox_E10 | 42 | 42 | 42 | 2 |
| TFIIS_M | 42 | 42 | 34 | 1 |
| mRNA_cap_enzyme | 41 | 41 | 41 | 1 |
| Pentapeptide_4 | 2 | 2 | 2 | 1 |
| Ank_2 | 97 | 75 | 45 | 0 |
| Pox_A32 | 83 | 1 | 0 | 0 |
| DUF4064 | 54 | 0 | 0 | 0 |
| ATPase_2 | 52 | 41 | 9 | 0 |
| DX | 49 | 0 | 0 | 0 |
| AAA_22 | 48 | 6 | 5 | 0 |
| Ank_4 | 46 | 44 | 0 | 0 |
| RNA_pol_A_bac | 42 | 8 | 0 | 0 |
| Kinase-like | 42 | 42 | 0 | 0 |
| PCNA_N | 42 | 42 | 0 | 0 |
| P12 | 42 | 42 | 34 | 0 |
| JIP_LZII | 42 | 10 | 0 | 0 |
| T5orf172 | 42 | 36 | 0 | 0 |
| DUF3169 | 42 | 0 | 0 | 0 |
| Spc7 | 42 | 34 | 0 | 0 |
| DUF3810 | 42 | 0 | 0 | 0 |
| HATPase_c | 42 | 42 | 42 | 0 |
| Baculo_LEF5_C | 42 | 8 | 0 | 0 |
| O-antigen_lig | 42 | 0 | 0 | 0 |
| DNA_gyraseB | 42 | 0 | 0 | 0 |
| RWD | 42 | 2 | 0 | 0 |
| SHS2_Rpb7-N | 42 | 42 | 8 | 0 |
| Amnionless | 42 | 22 | 0 | 0 |
| RNA_pol_Rpb1_6 | 42 | 10 | 0 | 0 |
| RIFIN | 42 | 1 | 0 | 0 |
| RNA_pol_Rpb2_5 | 42 | 10 | 0 | 0 |
| YibE_F | 41 | 0 | 0 | 0 |
| Bcl-2_3 | 41 | 0 | 0 | 0 |
| Colicin_V | 41 | 0 | 0 | 0 |
| Methyltransf_11 | 40 | 0 | 0 | 0 |
| DNA_pol3_a_NII | 40 | 0 | 0 | 0 |
| Nodulin_late | 38 | 31 | 2 | 0 |
| DUF4509 | 38 | 30 | 1 | 0 |
| Supplementary\_Table\_2\_ASFV\_PFAM\_domains, page 3 |  |  |  |  |
| NTP_transf_2 | 37 | 10 | 0 | 0 |
| Flavodoxin_3 | 36 | 0 | 0 | 0 |
| APH | 34 | 0 | 0 | 0 |
| RmuC | 34 | 0 | 0 | 0 |
| EzrA | 33 | 32 | 0 | 0 |
| ODV-E18 | 33 | 8 | 0 | 0 |
| Ank_5 | 31 | 29 | 9 | 0 |
| DUF1510 | 31 | 0 | 0 | 0 |
| Cation_ATPase_C | 30 | 0 | 0 | 0 |
| AAA_18 | 29 | 0 | 0 | 0 |
| Lectin_C | 26 | 25 | 0 | 0 |
| dNK | 25 | 1 | 0 | 0 |
| SRTM1 | 23 | 0 | 0 | 0 |
| AAA_17 | 20 | 20 | 0 | 0 |
| MGDG_synth | 19 | 0 | 0 | 0 |
| EB | 18 | 0 | 0 | 0 |
| Ank | 17 | 0 | 0 | 0 |
| DUF3394 | 17 | 1 | 0 | 0 |
| Actin_micro | 14 | 0 | 0 | 0 |
| HTH_26 | 14 | 0 | 0 | 0 |
| Prok-E2_B | 14 | 0 | 0 | 0 |
| Rep_2 | 13 | 1 | 0 | 0 |
| Ank_3 | 11 | 0 | 0 | 0 |
| PAN_4 | 11 | 2 | 0 | 0 |
| Stk19 | 10 | 0 | 0 | 0 |
| Methyltransf_23 | 9 | 2 | 0 | 0 |
| LapA_dom | 9 | 0 | 0 | 0 |
| AAA_14 | 9 | 0 | 0 | 0 |
| CTD | 8 | 0 | 0 | 0 |
| Renin_r | 7 | 0 | 0 | 0 |
| AAA_30 | 7 | 0 | 0 | 0 |
| DUF5381 | 7 | 0 | 0 | 0 |
| DUF302 | 6 | 6 | 0 | 0 |
| NB-ARC | 6 | 0 | 0 | 0 |
| DUF1189 | 6 | 0 | 0 | 0 |
| SirB | 6 | 0 | 0 | 0 |
| SieB | 6 | 0 | 0 | 0 |
| DUF2105 | 6 | 0 | 0 | 0 |
| KTI12 | 5 | 0 | 0 | 0 |
| DUF4153 | 5 | 0 | 0 | 0 |
| Synaptobrevin | 4 | 0 | 0 | 0 |
| TMEM154 | 4 | 0 | 0 | 0 |
| Pentapeptide_3 | 3 | 1 | 0 | 0 |
| Neurensin | 2 | 0 | 0 | 0 |
| Cpta_toxin | 2 | 1 | 0 | 0 |
| 3-HAO | 2 | 0 | 0 | 0 |
| DUF4512 | 1 | 0 | 0 | 0 |
| TIL | 1 | 0 | 0 | 0 |
| DUF587 | 1 | 0 | 0 | 0 |
| LRRNT_2 | 1 | 0 | 0 | 0 |
| DUF3772 | 1 | 0 | 0 | 0 |
| ABC2_membrane_5 | 1 | 0 | 0 | 0 |
| DUF3719 | 1 | 0 | 0 | 0 |
| Supplementary\_Table\_2\_ASFV\_PFAM\_domains, page 4 |  |  |  |  |
| CCDC158 | 1 | 0 | 0 | 0 |
| DUF5354 | 1 | 0 | 0 | 0 |
| DUF4904 | 1 | 0 | 0 | 0 |
| FUSC_2 | 1 | 0 | 0 | 0 |
| DNA_pol_B_palm | 1 | 0 | 0 | 0 |
| Sigma_reg_N | 1 | 0 | 0 | 0 |
| TRP | 1 | 0 | 0 | 0 |
| Imm39 | 1 | 0 | 0 | 0 |
| SelK_SelG | 1 | 0 | 0 | 0 |
| OAD_gamma | 1 | 0 | 0 | 0 |
| Clostridium_P47 | 1 | 0 | 0 | 0 |
| IncA | 1 | 0 | 0 | 0 |
| DUF4131 | 1 | 0 | 0 | 0 |
| Gemini_AL1_M | 1 | 0 | 0 | 0 |
| GP41 | 1 | 0 | 0 | 0 |
| Methyltransf_25 | 1 | 0 | 0 | 0 |
| Taxilin | 1 | 0 | 0 | 0 |
| DUF2937 | 1 | 0 | 0 | 0 |
| 7TM_GPCR_Sra | 1 | 0 | 0 | 0 |
| SpoIIIAH | 1 | 0 | 0 | 0 |
| DUF3749 | 1 | 0 | 0 | 0 |
| RNA_helicase | 1 | 0 | 0 | 0 |
| DUF4952 | 1 | 0 | 0 | 0 |
| Gly-zipper_OmpA | 1 | 0 | 0 | 0 |
| RE_SinI | 1 | 0 | 0 | 0 |
| Rft-1 | 1 | 0 | 0 | 0 |
| Arc_trans_TRASH | 1 | 0 | 0 | 0 |
| Orf78 | 1 | 0 | 0 | 0 |

## Readme document

### African Swine Fever Virus (ASFV) clustering tool

\# Below are instructions how to use the Docker image of the ASFV_classification tool to characterize ASFV genomes.

\# This tool is written and developed by Matthew Cotten and My Phan and described in the manuscript Masembe et al. (2019) “Increased resolution of African Swine Fever Virus genome patterns based on profile HMM protein domains”

### Please do not re-distribute this tool

The tool uses HMMR-3 from Sean R. Eddy and the HMMER development team (http://hmmer.org/) as well as a subset of the Pfam 31.0 library from (http://pfam.xfam.org/). We are extremely grateful to the developers of these tools for their open sharing of software and information.

Briefly, a fasta file of ASFV genome sequences is analyzed by the tool as follows: First, query sequences are screened for Pfam and custom MGF domains identified in the query ***Asfarviridae*** sequences at an e-value < 0.0001. The e-value, bit-score and position in the query sequence are gathered and the bit-scores are assembled into a matrix. Domains whose bit-scores vary across the set at variance >= to the variance argument are then used to cluster related ASFV sequences.

### To use the classification tool

1. Download a Docker from the website http://docs.docker.com/ Depending on your machine, you may download the Docker version for Windows, Linux, or Mac. If your current MacOS system is older than MacOS Yosemite 10.10.3, you may want to either upgrade your MacOS system. Alternatively, you can download the Docker Toolbox version at: http://docs.docker.com/docker-for-mac/docker-toolbox/
2. Follow the instructions on Docker webpage to install and test Docker.
3. Make a test_directory include your query fasta file in the directory and move to it: mkdir test_directory, cd test_directory
4. Within that directory, retrieve and load the docker image with the following command: docker pull matthewcotten/asfv_class_tool
5. To run the classification tool on sequences contained in the fasta file <asfv_sequences.fas=, use the following command: Note: Replace <path_to_test_directory= with the actual path to the directory, Replace <asfv_sequences.fas= with the actual name of the ASFV sequence file to be examined. /workdir is an actual directory within the docker image and this should not be confused with the test_directory. The <variance> argument sets the variance threshold for reporting domains in the final matrix used for clustering. We typically use 0.03 but investigators can experiment with alternate cutoffs. docker run -ti --rm -w /workdir -v <path_to_test_directory>:/workdir matthewcotten/asfv_class_tool Drop_hunt_ASFV_for_Docker4.py <asfv_sequences.fas= <variance=

### Output files

A. pdf of the clustermap.
B. A CSV table of domains identified with variance >= to the threshold that was set. For each domain identified the table lists the target genome and the location of the domain in the target genome (position, length and strand) and a variance flag (>= threshold variance in the set = high_variance, < threshold variance in the set = low_variance).

